# Application of the ANDROMEDA Software for Prediction of the Human Pharmacokinetics of Modern Anticancer Drugs

**DOI:** 10.1101/2023.03.18.533259

**Authors:** Urban Fagerholm, Sven Hellberg, Jonathan Alvarsson, Ola Spjuth

## Abstract

The ANDROMEDA toolkit for prediction of human clinical pharmacokinetics, based on machine learning, conformal prediction and a new physiologically-based pharmacokinetic model, was used to predict and characterize the human clinical pharmacokinetics of 12 small anticancer drugs marketed in 2021 and 2022 (molecular weight 355 to 1326 g/mol). The study is part of a series of software validations. A majority of clinical pharmacokinetic data was missing. ANDROMEDA successfully filled this gap. Most drugs were predicted/measured to have relatively complex pharmacokinetics, with limited passive permeability+efflux, high degree of plasma protein binding, significant gut-wall elimination and food interaction, biliary excretion and/or limited dissolution potential. Median, mean and maximum prediction errors for steady state volume of distribution, unbound fraction in plasma, blood-to-plasma concentration ratio, hepatic, renal and total clearance, fraction absorbed, oral bioavailability, half-life and degree of food interaction were 1.6-, 2.4- and 17-fold, respectively. Less than 3-fold errors were found for 78 % of predictions. Results are consistent with those obtained in previous validation studies and are better than with the best laboratory-based prediction methods, which validates ANDROMEDA for predictions of human clinical pharmacokinetics of modern small anticancer drugs with multi-mechanistical and challenging pharmacokinetics.

## Introduction

We have developed and validated an integrated *in silico*-based prediction software for human clinical pharmacokinetics (PK) – ANDROMEDA.^1–4^ The system is based on conformal prediction (CP), which is a methodology that sits on top of machine learning methods and produce valid levels of confidence,^5^ and a new human physiologically-based pharmacokinetic (PBPK) model.^1^ For a more extensive introduction to CP we refer to Alvarsson et al. (2021).^6^

ANDROMEDA has been used by our group for prediction of human clinical PK, including fraction absorbed (f_a_), oral bioavailability (F), half-life (t_½_), unbound fraction in plasma (f_u_), clearance (CL), steady-state volume of distribution (V_ss_), fraction excreted renally (f_e_) and food interaction.^2–4,7^

In a validation study (forward-looking predictions) with 65 chemicals of various classes, median and mean prediction errors for these PK-parameters were 1.3- to 2.7-fold and 1.4- to 4.8-fold, respectively.^4^ 59 % of predictions had prediction errors below 2-fold. In a validation study (forward-looking predictions) with pharmaceutical drugs, the predictive accuracy (Q^2^) for log V_ss_ was 0.65, and the average prediction error and % with <2-fold prediction error were 2.4-fold and 64 %, respectively.^2^ 69 % of compounds had an observed V_ss_ within the prediction interval at a 70 % confidence level, which empirically validated the software (it predicts at the chosen confidence level). Allometry was associated with higher mean error for the same set of compounds (2.9-fold).^2^ In another validation and comparative study with a benchmarking data set consisting of 24 physicochemically diverse compounds median prediction errors and Q^2^ for f_a_, F, f_u_, V_ss_, CL, f_e_, t_½_ and CL/F were 1.2- to 2.2-fold and 0.28 to 0.91, respectively.^3^ 82 % of predictions had maximum 3-fold error and the maximum prediction error was 15-fold (for a compound with 1 % F).^3^ In 76 and 12 % of comparisons of predictive accuracy, ANDROMEDA was superior to or on par with laboratory methods, respectively.^3^ In addition to the validation studies, ANDROMEDA predictions were approved by the German authority BfArM for use and main preclinical PK-source in clinical trial applications.

New drugs on the market give an opportunity to evaluate the validity and accuracies of prediction systems and software for modern compounds (with higher molecular weight (MW) and (anticipated) multi-mechanistical and challenging PK). Anticancer drugs make up a significant portion (ca ¼) of all new small drugs, and studies demonstrating the ability of methods to predict their PK are lacking.

The main objective of this study was to validate ANDROMEDA for prediction of the human clinical PK of small modern anticancer drugs. Another aim was to characterize the PK of these drugs.

## Methods

### ANDROMEDA PK-Prediction Software

For a description of ANDROMEDA (including underlying CP models, algorithms, molecular descriptors, parameters and performance) see Fagerholm et al. 2022.^2,4^ The software predicts 30 human PK-parameters, including V_ss_, f_u_, blood-to-plasma concentration ratio (C_bl_/C_pl_), CL_H_, renal C_L_ (C_LR_), f_e_, biliary CL (CL_bile_), CL, CL/F, oral V_ss_ (V_ss_/F), fraction escaping gut-wall extraction (F_gut_), f_a_, F, t_½_, and MDR-1-, BCRP-, MRP2-, CYP2C9-, CYP2D6- and CYP3A4-specificities (parameters of main interest in this study).^1–4^ ANDROMEDA-data can also be used to predict the degree of food interaction (ratio between area under the plasma concentration *vs* time profile in fed (food with high fat content) and fasted states).^7^

Prediction accuracy/errors were estimated as predicted/observed or observed/predicted values (ratios≥1 were selected) and correct classification (substrate *vs* non-substrate for CYPs and transporters).

ANDROMEDA is mainly applicable for compounds with MW 100 to 700 g/mol and for non-saturated conditions (not a high doses). There are groups of compounds for which the *in silico* models do not work, including metals and quaternary amines, and have limited use, for example, hydrolysis sensitive compounds and drugs binding covalently and/or to DNA (some anticancer drugs bind to DNA).

None of the new small drugs in the study was included in the training sets of the used CP models. Thus, every prediction was a forward-looking prediction where each compound was unknown to the models.

### Compound Selection and Clinical PK-Data

Small anticancer compounds marketed in 2021 and 2022 were explored and selected at U.S. Food and Drug Administration (FDA). Only hydrolysis-insensitive drugs with a maximum MW of 1500 g/mol were selected for the study. This resulted in a selection of 12 compounds with MW 355 to 1326 g/mol (6 compounds with a MW exceeding 500 g/mol). One of the compounds is dosed intravenously (pafolacianine), whereas the remainder of drugs are administered orally. PK-data were mainly selected from FDA-label documents (Clinical Pharmacology sections) of each drug.

## Results

The amount of clinical PK-data for the compounds was limited (more than 50 % was missing) – V_ss_ (n=1), f_u_ (n=12), C_bl_/C_pl_ (n=4), CL (n=1), CL_H_ (n=1), CL_R_ (n=1), CL/F (n=11), f_a_ (n=6), F (n=2), t_½_ (n=12), substrate specificity for CYPs 2C9, 2D6 and/or 3A4 (n=12), substrate specificity for MDR-1, BCRP and/or MRP2 (n=9), bile excretion (n=0), CL_bile_ (n=0), F_gut_ (n=0) and food interaction (n=11). Ranges for V_ss_, f_u_, C_bl_/C_pl_, CL, CL_H_, CL_R_, CL/F, f_a_, F, t_½_ and food interaction were 0.24 L/kg, <0.003-0.11, 0.56 to 0.76, 477 mL/min, 386 mL/min, <91 mL/min, 13-2300 mL/min, >0.62-near complete, 0.37-0.72, 0.44-111 h and 0.38-2.00-fold, respectively.

ANDROMEDA was capable of predicting the human clinical PK for all the selected compounds. Prediction results are shown in Table 1. Median, mean and maximum prediction errors for V_ss_, f_u_, C_bl_/C_pl_, CL, CL_H_, CL_R_, CL/F, f_a_, F, t_½_ and food interaction were 1.01 to 3.78-, 1.01 to 2.59- and 1.01 to 17-fold, respectively. The overall median and mean prediction errors were 1.56- and 2.39-fold, respectively, and 78 % of predictions had a maximum 3-fold error. Incorrect classifications for CYPs (2C9, 2D6 and/or 3A4) and transporters (MDR-1, BCRP and/or MRP2) were found for 15 % of the compounds, respectively.

**Table 1.** Observed and predicted human pharmacokinetics of 12 small anticancer drugs marketed in 2021 and 2022.

|  | MW | Route of |  | F | fa | t½ | Vss | fu | Cbl/Cpl | CL | CLH | CLR | CL/F | fe | Food interaction |
| --- | --- | --- | --- | --- | --- | --- | --- | --- | --- | --- | --- | --- | --- | --- | --- |
|  | [g/mol] | administration |  | fraction | fraction | [h] | [L/kg] | fraction | fraction | [mL/min] | [mL/min] | [mL/min] | [mL/min] | fraction | [-fold] |
| Adagrasib | 604 | po | Observed | - | >=0,81 | 23 | - | 0,020 | - | - | - | - | 616,7 | - | 1,00 |
|  |  |  | Predicted | - | 0,78 | 12 | - | 0,023 | - | - | - | - | 1242,4 | - | 1,36 |
|  |  |  | Error | - | 1,15 | 1,92 | - | 1,14 | - | - | - | - | 2,01 | - | 1,36 |
| Asciminib | 450 | po | Observed | - | 0,98 | 5,5 | - | 0,03 | - | - | - | - | 112 | - | 0,38 |
|  |  |  | Predicted | - | 0,82 | 3,1 | - | 0,076 | - | - | - | - | 567 | - | 1,07 |
|  |  |  | Error | - | 1,19 | 1,76 | - | 2,53 | - | - | - | - | 5,07 | - | 2,82 |
| Futibatinib | 418 | po | Observed | - | - | 2,9 | - | 0,050 | - | - | - | - | 333,3 | - | 0,89 |
|  |  |  | Predicted | - | - | 3,3 | - | 0,202 | - | - | - | - | 496,6 | - | 0,57 |
|  |  |  | Error | - | - | 1,14 | - | 4,04 | - | - | - | - | 1,49 | - | 1,56 |
| Infigratinib | 560 | po | Observed | - | - | 34 | - | 0,032 | - | - | - | - | 552 | 0,02 of F | 2,00 |
|  |  |  | Predicted | - | - | 8,2 | - | 0,012 | - | - | - | - | 332 | 0,01 | 1,23 |
|  |  |  | Error | - | - | 4,09 | - | 2,64 | - | - | - | - | 1,66 | - | 1,63 |
| Mobocertinib | 586 | po | Observed | 0,37 | 0,94 | 18 | - | 0,007 | 0,76 | - | - | - | 2300 | 0,25 | 1,00 |
|  |  |  | Predicted | 0,49 | 1,00 | 4,6 | - | 0,029 | 0,69 | - | - | - | 746 | 0,0002 | 1,30 |
|  |  |  | Error | 1,32 | 1,06 | 3,96 | - | 4,14 | 1,10 | - | - | - | 3,08 | - | 1,30 |
| Olutasidenib | 355 | po | Observed | - | >=0,62 | 67 | - | 0,070 | - | - | - | - | 66,7 | - | 1,83 |
|  |  |  | Predicted | - | 0,88 | 23 | - | 0,023 | - | - | - | - | 57,9 | - | 0,96 |
|  |  |  | Error | - | 1,09 | 2,88 | - | 3,070 | - | - | - | - | 1,15 | - | 1,91 |
| Pacritinib | 473 | po | Observed | - | Ca 1 | 28 | - | 0,012 | - | - | - | - | 34,8 | - | 1,00 |
|  |  |  | Predicted | - | 0,69 | 7,3 | - | 0,028 | - | - | - | - | 581,9 | - | 1,64 |
|  |  |  | Error | - | 1,43 | 3,79 | - | 2,29 | - | - | - | - | 16,71 | - | 1,64 |
| Pafolacianine | 1326 | iv | Observed | - | - | 0,44 | 0,24 | 0,06 | 0,56 | 477 | 386 | <=91 | - | Max 0,19 | - |
|  |  |  | Predicted | - | - | 1,0 | 0,45 | 0,06 | 0,64 | 450 | 389 | 7,7 | - | 0,02 | - |
|  |  |  | Error | - | - | 2,27 | 1,88 | 1,02 | 1,14 | 1,06 | 1,01 | <12 | - | <9,5 | - |
| Sotorasib | 561 | po | Observed | - | - | 5,3 | - | 0,11 | - | - | - | - | 437 | 0,01 of F | 1,25 |
|  |  |  | Predicted | - | - | 4,2 | - | 0,01 | - | - | - | - | 537 | 0,0 | 1,50 |
|  |  |  | Error | - | - | 1,26 | - | 11,0 | - | - | - | - | 1,23 | - | 1,20 |
| Tepotinib | 493 | po | Observed | 0,72 | >0,72 | 32 | - | 0,02 | - | - | - | - | 397 | 0,07 of F | 1,60 |
|  |  |  | Predicted | 0,57 | 0,89 | 15 | - | 0,025 | - | - | - | - | 280 | 0,03 | 1,59 |
|  |  |  | Error | 1,25 | 1,04 | 2,19 | - | 1,25 | - | - | - | - | 1,42 | - | 1,01 |
| Tivozanib | 455 | po | Observed | - | - | 111 | - | <0,01 | 0,56 | - | - | - | 12,5 | 0 of F | 1,00 |
|  |  |  | Predicted | - | - | 30 | - | 0,049 | 0,60 | - | - | - | 44,7 | 0,0 | 1,22 |
|  |  |  | Error | - | - | 3,66 | - | - | 1,07 | - | - | - | 3,58 | - | 1,22 |
| Umbralisib | 572 | po | Observed | - | - | 91 | - | <0,003 | 0,60 | - | - | - | 258 | 0,0002 of F | 1,61 |
|  |  |  | Predicted | - | - | 32 | - | 0,004 | 0,61 | - | - | - | 63 | 0,0001 | 1,70 |
|  |  |  | Error | - | - | 2,81 | - | - | 1,02 | - | - | - | 4,13 | - | 1,06 |

|  | MW | Route of |  | Bile exc | CYPs | Transporters |
| --- | --- | --- | --- | --- | --- | --- |
|  | [g/mol] | administration |  |  | 2C9,2D6,3A4 | MDR1,BCRP,MRP2 |
| Adagrasib | 604 | po | Observed | - | Yes | Yes? |
|  |  |  | Predicted | Yes | Yes | Yes |
|  |  |  | Error | - | Correct | Correct? |
| Asciminib | 450 | po | Observed | - | Yes | Yes |
|  |  |  | Predicted | Ud | Yes | Yes |
|  |  |  | Error | - | Correct | Correct |
| Futibatinib | 418 | po | Observed | - | Yes | Yes |
|  |  |  | Predicted | Ud | Yes? | Yes |
|  |  |  | Error | - | Correct? | Correct |
| Infigratinib | 560 | po | Observed | - | Yes | Yes |
|  |  |  | Predicted | No | Yes | Yes |
|  |  |  | Error | - | Correct | Correct |
| Mobocertinib | 586 | po | Observed | - | Yes | Yes |
|  |  |  | Predicted | Yes | Yes | Yes |
|  |  |  | Error | - | Correct | Correct |
| Olutasidenib | 355 | po | Observed | - | Yes | ? |
|  |  |  | Predicted | No | Yes | Yes |
|  |  |  | Error | - | Correct | - |
| Pacritinib | 473 | po | Observed | - | Yes | No |
|  |  |  | Predicted | No | Yes | Yes |
|  |  |  | Error | - | Correct | Incorrect |
| Pafolacianine | 1326 | iv | Observed | - | No | - |
|  |  |  | Predicted | Yes | Yes | - |
|  |  |  | Error | - | Incorrect | - |
| Sotorasib | 561 | po | Observed | - | Yes | - |
|  |  |  | Predicted | Yes | Yes | - |
|  |  |  | Error | - | Correct | - |
| Tepotinib | 493 | po | Observed | - | Yes | Yes |
|  |  |  | Predicted | No | Yes | Yes |
|  |  |  | Error | - | Correct | Correct |
| Tivozanib | 455 | po | Observed | - | Yes | No |
|  |  |  | Predicted | No | Yes | Yes |
|  |  |  | Error | - | Correct | Incorrect |
| Umbralisib | 572 | po | Observed | - | Yes | - |
|  |  |  | Predicted | No | Yes | - |
|  |  |  | Error | - | Correct | - |

The PK of the largest compound (pafolacianine; MW=1326 g/mol) was generally well predicted.

According to predictions, no compound is significantly (≥20%) eliminated via the renal route, 75 % of compounds are significantly (≥20%) degraded in the gut-wall, 33 % of compounds are excreted via bile, 25 % of compounds have low/moderate passive intestinal permeability with efflux, 50 % of compounds have limited gastrointestinal dissolution/solubility potential, no compound has a metabolic stability that is normally problematic to quantify with human hepatocytes (approximately <500-1000 mL/min), 50 % of compounds have very low f_u_ (≤0.02) (also 50 % according to measurements) and 58 % of compounds have significant (but not extensive) food interaction (Figure 1).

**Figure 1.**
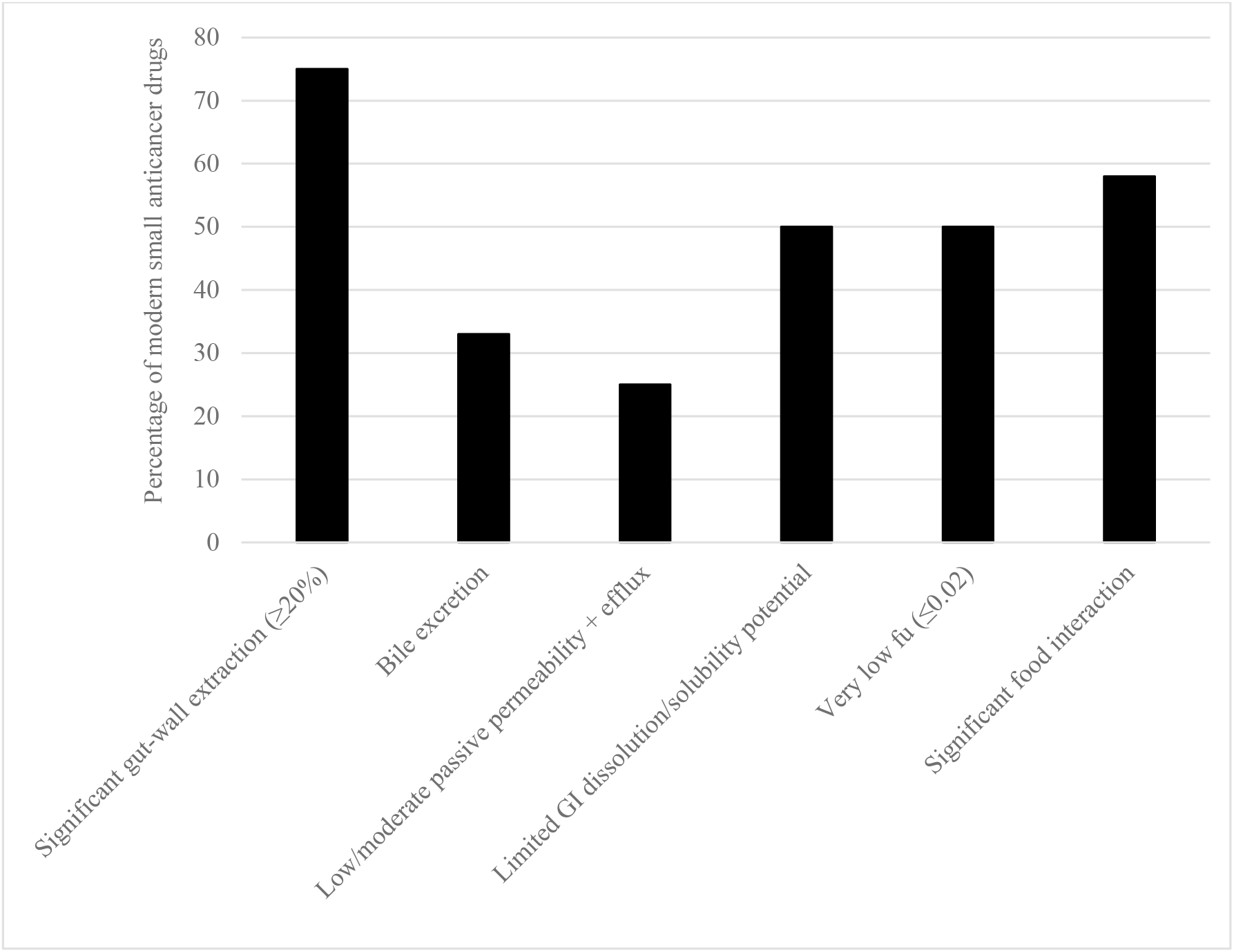
The percentage of modern small anticancer drugs with certain predicted PK characteristics.

## Discussion

This study further validates ANDROMEDA for prediction of human clinical PK. Results are consistent with those obtained in earlier validation studies with the software.^2–4,7^

The total 1.6-fold median prediction error is similar to or lower than the median lab variability for f_u_ and intrinsic metabolic CL (CL_int_).^8–10^

Apparently, the results are equally good as or better than those obtained with best of comparable laboratory methods.^3,11,12^ The mean prediction errors for CL/F (3.8-fold) and t_½_ (2.6-fold) were similar to or better than corresponding absolute average fold errors obtained with the best of animal data-based models (4.2- and 2.4-fold, respectively) and *in vitro* data-based methods (7.8- and 2.5-fold, respectively) in the PhRMA CPCDC initiative (consisting of a number of major international pharmaceutical companies).^12^ An approximate 4-fold mean prediction error for CL/F was also produced in the laboratory by Miljković et al.^11^ In contrast to the laboratory methods (with 1/4 and 2/3 excluded/non-quantifiable compounds with animal and *in vitro*-models, respectively) ANDROMEDA produced complete sets of predictions for all compounds.^12^ Thus, the software seems to have a superior application domain compared to laboratory methods.

The greatest prediction errors (11- and 17-fold) were found for compounds with high degree of plasma protein binding. A recent evaluation showed that there is also considerable lab variability for highly bound compounds – maximum 185-fold and >5-fold for significant portion of compounds with average f_u_≤0.01^8^ – indicating greater uncertainty of laboratory method-based predictions as well. Maximum prediction errors of up to million-fold have been found for laboratory methods.^1,3^

Deviating substrate specificities for CYPs and transporters occur with both our methodology (15 % incorrect classification predictions) and lab methods (10 and 34 % inconsistency between studies for CYP3A4 and MDR-1, respectively).^13^

It is worth noting the limited amount of PK-information in FDA-label documents for the new small anticancer drugs (most data missing) and the ability of ANDROMEDA to add missing data and knowledge.

Significant gut-wall extraction, bile excretion, limited permeability, limited solubility/dissolution very low f_u_ and/or significant food interaction was predicted for many of the new anticancer drugs, which demonstrates the requirement of methods to predict these processes for this class of small drugs *in vivo* in man. ANDROMEDA has this capacity.

Limitations with ANDROMEDA include the limited molecular weight range (mainly applicable for compounds with MW 100 to 700 g/mol), and lack of general capability to predict dose- and time-dependencies and capability to predict V_ss_ and overall PK for compounds that bind/intercalate significantly to DNA. For example, the V_ss_ of DNA-binding/intercalating anticancer compounds doxorubicin, epirubicin, idarubicin, mitoxantrone and vinblastine is in the 10-50 L/kg range. For such compounds, human V_ss_ is probably predicted better from animal data.

Prediction of V_ss_ and t_½_ for biliary excreted compounds with enterohepatic circulation is challenging both for ANDROMEDA and laboratory methods. Despite this, ANDROMEDA predicted the t_½_ of anticancer drugs with anticipated/predicted bile excretion and enterohepatic circulation well.

In conclusion, the results are consistent with those obtained in previous validation studies and validates ANDROMEDA for predictions of human clinical PK of modern small anticancer drugs, which to a great extent demonstrate quite complex PK-characteristics with absorption obstacles (limited permeability and/or solubility/dissolution) and/or multiple elimination pathways (hepatic metabolism, gut-wall metabolism and renal and biliary excretion).

## References

1. Fagerholm U, Hellberg S, Spjuth O. Advances in predictions of oral bioavailability of candidate drugs in man with new machine learning methodology. Molecules 2021;26:2572.

2. Fagerholm U, Hellberg S, Alvarsson J. Spjuth O. *In silico* prediction of volume of distribution of drugs in man using conformal prediction performs on par with animal data-based models. Xenobiot. 2021;51(12):1366–1371.

3. Fagerholm U, Hellberg S, Alvarsson J, Spjuth O. *In silico* prediction of human clinical pharmacokinetics with ANDROMEDA by Prosilico: Predictions for an established benchmarking data set, a modern small drug data set, and a comparison with laboratory methods. Altern Lab Anim. 2023;51:39–54.

4. Fagerholm U, Hellberg S, Alvarsson J, Spjuth O. *In silico* predictions of the human pharmacokinetics/toxicokinetics of 65 chemicals from various classes using conformal prediction methodology. Xenobiot. March 2022

5. Vovk V, Gammerman A, Shafer G. Algorithmic learning in a random world. Springer Science & Business Media, 2005.

6. Alvarsson J, Arvidsson McShane S, Norinder U. Spjuth O. Predicting with confidence: using conformal prediction in drug discovery. J Pharm Sci. 2021;110:42–49.

7. Fagerholm U, Hellberg S, Alvarsson J, Spjuth O. Predicting the influence of fat food intake on the absorption and systemic exposure of modern small drugs using ANDROMEDA by Prosilico software. bioRxiv, Dec 2022.

8. Fagerholm U, Spjuth O, Hellberg S. Comparison between lab variability and *in silico* prediction errors for the unbound fraction of drugs in human plasma. Xenobiot. 2021;51:1095–1100.

9. Sohlenius-Sternbeck A-K, Afzelius L, Prusis P, Neelissen J, Hoogstraate J, Johansson J, Floby E, Bengtsson A, Gissberg O, Sternbeck J, Petersson C. Evaluation of the human prediction of clearance from hepatocyte and microsome intrinsic clearance for 52 drug compounds. Xenobiot. 2010;40:637–649.

10. Yamagata T, Zanelli U, Gallemann D, Perrin D, Dolgos H, Petersson C. Comparison of methods for the prediction of human clearance from hepatocyte intrinsic clearance for a set of reference compounds and an external evaluation set. Xenobiot. 2017;47:741–751.

11. Miljković F, Martinsson A, Obrezanova O, Williamson B, Johnson M, Sykes A. Machine learning models for human *in vivo* pharmacokinetic parameters with in-house validation. Mol Pharmaceut. November 10, 2021.

12. Poulin P, Jones RDO, Jones HM, Gibson CT, Rowland M, Chien JY, Ring BJ, Adkinson KK, Ku S, Vuppugalla R, Marathe P, Fischer O, Dutta S, Sinha VK, Björnsson T, Lavé T. Yates JW. PHRMA CPCDC initiative on predictive models of human pharmacokinetics, Part 5: Prediction of plasma concentration– time profiles in human by using the physiologically-based pharmacokinetic modeling approach. J Pharm Sci. 2011;100:4127–4157.

13. Fagerholm U. Comparing *in silico* and *in vitro* methods for classification of BCS II and CYP3A4 and MDR-1 substrate specificity. bioRxiv, Dec 2022.

